# *Psidium guajava* in the Galapagos Islands: population genetics and history of an invasive species

**DOI:** 10.1101/402693

**Authors:** Diego Urquía, Bernardo Gutiérrez, Gabriela Pozo, María José Pozo, Analía Espín, María de Lourdes Torres

**Affiliations:** Universidad San Francisco de Quito (USFQ). Colegio de Ciencias Biológicas y Ambientales, Laboratorio de Biotecnología Vegetal. Campus Cumbayá. Quito, Ecuador.; Galapagos Science Center, Universidad San Francisco de Quito and University of North Carolina at Chapel Hill, San Cristobal, Galapagos, Ecuador; Department of Zoology. University of Oxford. South Parks Road, Oxford OX1 3SY, United Kingdom.

## Abstract

The threat of invasive plant species in island populations prompts the need to better understand their population genetics and dynamics. In the Galapagos islands, this is exemplified by the introduced guava (*Psidium guajava*), considered one of the greatest threats to the local biodiversity due to its effective spread in the archipelago and its ability to outcompete endemic species. To better understand its history and genetics, we analyzed individuals from three inhabited islands in the Galapagos archipelago with 11 SSR markers. Our results reveal similar genetic diversity between islands, suggestive of gene flow between them. Populations appear to be distinct between the islands of San Cristobal and Isabela, with the population of Santa Cruz being composed as a mixture from both. Additional evidence for genetic bottlenecks and the inference of introduction events suggests an original introduction of the species in San Cristobal, from where it was later introduced to Isabela, and finally into Santa Cruz. Alternatively, an independent introduction event for Isabela is also possible. These results are contrasted with the historical record, providing a first overview of the history of *P. guajava* in the Galapagos islands and its current population dynamics.

## Introduction

Invasive plant species are a serious threat to natural ecosystems and biological diversity [1, 2]. Studies have shown that they can cause a reduction of the local biodiversity, modify the compositions of resident communities, change nutrient cycling processes and are attributed an overall alteration of ecosystem properties [3, 4, 5]. Once an invasive plant species has been introduced into a new ecosystem, it has the capacity to spread and consolidate, and potentially cause severe repercussions for native flora and fauna, even leading to their extinction [1]. Islands are particularly vulnerable to invasive species because of their isolation from mainland and generally low biodiversity [6]. This susceptibility is most evident for endemic species, given the unique selective forces that island ecosystems can exert on resident organisms, fine-tuning their adaptive profile to make them highly specialized for their niches [6]. These species are usually less genetically diverse, have weak crossing barriers and unspecialized pollinators, all of which make them more susceptible to invaders [7].

The Galapagos Islands are an example of an isolated ecosystem vulnerable to invasive plant species due to their unique biodiversity and high biological endemism [8, 9]. Aside from being an isolated archipelago, the Galapagos were only recently inhabited by humans (1830s), making them an ideal setting to study the introduction of invasive species [10]. Since their discovery in 1535, a total of 821 alien plant species have been introduced into the islands [9, 11]. History has shown that endemic species rapidly become extinct when alien species are introduced, and invasive species have been formally recognized as a serious threat to the Galapagos Islands’ ecosystems [10, 12]. There have been few studies that explore the impact of invasive plants in the Galapagos; one such study reported a strong negative effect of the invasive plant *Rubus niveus* on the species richness of the *Scalesia* forests [5].

The common guava (*Psidium guajava*) is an invasive plant species in the Galapagos Islands. It is commonly cultivated in South Asia, Central and South America, North America and Australia for its edible fruit[13]. It is believed that *P. guajava* originated in Central and South America [13] and was introduced to the Galapagos Islands in the late 19th century [12, 14]. It is now recognized as one of the 37 highly invasive plants in the archipelago, having settled in non-cultivated areas since the early 1900s [6, 12]. Currently, *P. guajava* has a widespread distribution in the Galapagos, growing in disturbed areas as well as in natural forests in Isabela, Santa Cruz, San Cristobal and Floreana, which are also the islands of the archipelago that host human populations [12, 15]. In addition, *P. guajava* could pose a threat to closely related species such as the endemic congeneric guayabillo (*Psidium galapageium)* [16]. In the Galapagos, both species partially share their spatial distribution, potentially making them direct competitors and candidates for interspecific hybridization [16]. Extinction of insular plant species via hybridization between congeners has been well-established: when it occurs, it can reduce a native species’ population growth by altering its interactions with other species and affecting its reproductive and competitive success, eventually leading to its extinction [17, 18]. This is exemplified by the ongoing hybridization in Socorro Island (Mexico) between native species *Psidium socorrense* and its congener *P*. sp. aff. *Sartorianum* which is resulting in the local extinction of the native species at the southern slope of the island [18].

Genetic diversity and population structure are important factors when examining the origins, introduction history and invasion path of an alien species, including the number of introduction events [19]. As *P. guajava* affects the archipelago’s ecosystems and could be a threat to its endemic congener, the guayabillo, it is important to understand its genetic profile to adequately model its invasive dynamics. Here we analyze the genetic diversity and population structure of *P. guajava* populations on the Isabela, Santa Cruz and San Cristobal islands in the Galapagos (Ecuador) to understand the history and current status of this species in the archipelago.

## Methods

### Sampling and DNA Extraction

A total of 269 *P. guajava* individuals were sampled from selected locations of three islands in the Galapagos archipelago: 94 samples were obtained from San Cristobal (shortened SCY), 80 from Santa Cruz (shortened SCZ) and 95 from Isabela (shortened ISA). The sampled sites encompass the area on each island that is accessible by main or secondary roads, and focus on areas where human settlements or agricultural activities are widespread (due to the historical link between *P. guajava* and human settlements). Two to five young leaves were collected from each individual and transported to the Molecular Biology and Microbiology Laboratory at the Galapagos Science Center in San Cristobal, where they were stored at −20°C. Collection sites were georeferenced using a Garmin E-Trex Legend HCx GPS system (Garmin International Inc., USA).

Total genomic DNA was extracted from leaves using the CTAB protocol described by Saghai-Maroof *et al*. [20]. DNA concentration and quality were assessed using a Nanodrop 1000 Spectrophotometer (Thermo Scientific, USA). DNA was then transported to the Plant Biotechnology Laboratory at Universidad San Francisco de Quito for further processing.

### Sample preparation and genotyping

Thirteen SSR markers for *P. guajava* developed by Risterucci *et al*. were used for this study [21]. For the PCR amplification of microsatellite loci, annealing temperatures were optimized for each set of primers; cycling conditions were 15 min at 95°C, followed by 30 to 40 cycles of 30 sec at 94°C, 90 sec at the standardized annealing temperature, 60 sec at 72°C, and a final elongation step of 5 min at 72°C. PCR products were labeled with one of four fluorescent dyes (6-FAM, VIC, PET or NED) using universal primers in a three-primer system described by Blacket, *et al*. [22]. Labeled PCR products were commercially genotyped by Macrogen (Seoul, Korea) on an ABI 3130 Genetic Analyzer (ThermoFisher Scientific, USA) using 500LIZ as a size standard. Genotyping results were analyzed using the GeneMarker software v. 2.4.0 (Softgenetics, State College, PA, USA).

### Data analysis

#### Genetic diversity and population genetics

Maps of the georeferenced sampling regions on each island were drawn using ARCGIS Desktop 10.2 (Environmental Systems Research Institute, CA, USA). For each SSR locus analyzed, average number of alleles (A), observed heterozygosity (H_O_), expected heterozygosity (H_E_) and F-statistics were estimated on the *hierfstat* package [23] as implemented in R [24]. Private alleles (PA) were determined using the *poppr* package [25] for R. A Linkage Disequilibrium (LD) tests and an analysis of molecular variance (AMOVA) to test the genetic differentiation between islands and between regions within each island were conducted using Arlequin 3.5 [26]. The *adegenet* package [27] was used to evaluate the distribution of fixation indices on each island as an indicator of the proportion of inbreeding in each population. The same package was also used to perform a non-parametric Monte-Carlo test to evaluate the differences in H_E_ between islands. Finally, Hardy-Weinberg Equilibrium for each marker was tested with the *pegas* package [28] for R. Bonferroni corrections for multiple paired comparisons were applied for the LD and Monte-Carlo tests.

#### Evaluation of population structure

Pairwise F_ST_ genetic distances [29] were calculated with the *hierfstat* package, and a principal coordinate analysis (PCoA) was plotted with the *ggord* package [30] to quantify the differentiation between islands (or regions within islands) and visualize the genetic structure. In addition, STRUCTURE 2.3.4 [31] was used to infer population structure using a Bayesian individual-based clustering approach. The program was run with an admixture model, using individual sampling islands as a prior. The potential number of genetic clusters (*K*) was evaluated between 1 and 10, with 10 independent runs performed for each *K* value. 1,000,000 Markov Chain Monte Carlo (MCMC) steps were used, with a 100,000-step burn-in period. The optimum value of *K* was evaluated using the Evanno method [32] as implemented in Structure Harvester [33], and individual membership coefficients were summarized from independent runs with the program CLUMPP [34]. The final STRUCTURE graph was plotted using the *pophelper* package [35] implemented in R. Following the same procedure, STRUCTURE plots for each island individually were also obtained (where no geographical information was included as a prior).

#### Population history of *P. guajava* in the Galapagos Islands

Approximate Bayesian Computation (ABC) analyses were run on DIYABC 2.0 [36] to infer the colonization patterns and introduction history of *P. guajava* in the Galapagos Islands. A total of 14 different colonization scenarios (S1 File) were compared through 7,000,000 simulations following a stepwise mutation model (SMM) for microsatellite loci. Posterior probabilities of each scenario were computed using the logistic approach in DIYABC 2.0. The times of divergence (t1 and t2) of populations (in generations) according to the best supported scenarios were also estimated. To better understand these patterns and confirm possible introduction events, the potential of population bottlenecks in every island was inferred through the BOTTLENECK software, following both, the SMM and TPM models [37]. Heterozygosity excess or deficiencies were tested through a Wilcoxon Sign-Rank test.

## Results

### Genetic diversity of *P. guajava* in the Galapagos Islands

Genetic information for *P. guajava* individuals from three islands in the archipelago was successfully obtained for 11 of the 13 nuclear SSR markers tested (S1 Table). The number of total alleles differs markedly between the three islands, with the Isabela population containing the highest number of alleles (40 alleles) and San Cristobal the lowest (25 alleles). However, when counting the frequent alleles exclusively (defined here as alleles which occur at a frequency >0.05; see [38]), we found that the numbers were very similar (20-22 frequent alleles) among the three islands (Table 1). The number of exclusive alleles for each island differs greatly, ranging from 2 in San Cristobal to 15 in Isabela. The latter is also the only population that displays frequent exclusive alleles, defined as alleles that occur at frequencies higher than 5% in its population and are not found in any other island (Table 1).

**Table 1.** Genetic diversity information of the analyzed *Psidium guajava* populations from Isabela, Santa Cruz and San Cristobal islands: Number of individuals genotyped from each island (N), number of alleles found (A), number of private alleles (PA), observed heterozygosity (H_O_), expected heterozygosity/gene diversity (H_E_) and inbreeding coefficient (F_IS_). Overall results along the three islands are also shown.

| Island | N | A* | PA* | $H_O^a$ | $H_E^a$ | $F_{IS}$ |
| --- | --- | --- | --- | --- | --- | --- |
| Isabela | 95 | 40 (22) | 15 (4) | 0.106 | 0.284 | 0.621 |
| Santa Cruz | 80 | 35 (21) | 8 (0) | 0.169 | 0.365 | 0.539 |
| San Cristobal | 94 | 25 (20) | 2 (0) | 0.213 | 0.326 | 0.341 |
| **Overall** | 269 | 52 (21) | - | 0.163 | 0.356 | 0.505 |

\* Values between brackets are the number of alleles or private alleles with a frequency $>0.05$ within the corresponding island population.
<sup>a</sup>indicates average across the 11 SSRs analyzed.

The expected heterozygosity (H_E_) estimates show a greater genetic diversity in Santa Cruz (H_E_ = 0.365; SD = 0.202) and San Cristobal (H_E_ = 0.326, SD = 0.230), with the lowest value found in Isabela (H_E_ = 0.284, SD = 0.133) (Table 1). These differences in H_E_ between islands were not significant between Isabela and San Cristobal (after a Bonferroni correction, *p* = 0.037), but were significant between Santa Cruz and both Isabela (*p* = 0.001) and San Cristobal (*p* = 0.005). Linkage disequilibrium between markers was not found for Santa Cruz (and was spuriously found in one pair of markers in San Cristobal) but appears to be relatively common in Isabela (S2 Table). This location-dependent linkage may suggest the effect of some evolutionary forces in the Isabela population rather than linkage due to genomic proximity between markers, therefore permitting the assumption of independent segregation for all posterior analyses. Hardy-Weinberg Equilibrium was also tested for all markers, indicating significant disequilibrium for most markers in all islands (S3 Table).

The degree of inbreeding in each of the three islands, explored through the F_IS_ statistic, revealed values ranging from 0.341 to 0.621 (Table 1). The overall inbreeding coefficient for all three islands appears higher than the values for each individual island, and a visual inspection of the distribution of inbreeding coefficients in the archipelago shows a skewed, non-normal distribution (S1 Fig), suggesting that some inbreeding could occur in a proportion of the sampled individuals.

### Population structure

The genetic differentiation of *P. guajava* populations across the archipelago is relatively low. Only 13% of the variation was explained by the separation between islands. When analyzing the two most divergent populations, Isabela and San Cristobal, the molecular variance explained by populations only rose to 20%, suggesting that a large degree of genetic similarities is shared between islands (Table 2). The analysis of differentiation between sampling locations within each island reveals even lower proportions of the molecular variance embedded in this level, suggesting no differentiation within each island (S4 Table). However, it should be noted that, despite this low differentiation between islands and regions within islands, gene flow appears to be defined by proximity in an isolation-by-distance fashion, as the pairwise F_ST_ genetic distances between regions within each island or region reached considerably high values (S5 Table – S7 Table).

**Table 2.** Results of the analysis of molecular variance (AMOVA) performed for the *Psidium guajava* populations of Isabela, Santa Cruz and San Cristobal islands, and over the Isabela and San Cristobal populations, excluding Santa Cruz. Missing data was ignored for the AMOVA calculations.

| Source of variation | Three islands |  | Isabela & San Cristobal |  |
| --- | --- | --- | --- | --- |
|  | % of variation | *p*-value | % of variation | *p*-value |
| Between Islands | 13.17 | 0.001 | 20.05 | 0.001 |
| Between samples within Island | 44.24 | 0.001 | 39.24 | 0.001 |
| Within samples | 42.58 | 0.001 | 40.71 | 0.001 |

To further explore the degree of genetic structure in these populations, different approaches were considered. A Euclidean-based classification of genetic distances between individuals shows minor differentiation between islands, even when the top two eigenvectors explain over 60% of the total variance (Fig. 2). This approach shows a greater differentiation between the populations from Isabela and San Cristobal, the two furthest apart islands in our study. Furthermore, a Bayesian-based estimation of the number of clusters (*K*) that best explain the population structure in our data set suggested an optimal number of two lineages which cluster San Cristobal and Isabela under different groups (Fig. 3, orange and green respectively), with the population of Santa Cruz being composed of a combination of both lineages, with a clear dominance of the San Cristobal lineage (Fig. 3). A similar analysis of the samples within each island reveals no defined structure for their respective *P. guajava* populations (S2 Fig – S4 Fig). The quantification of the differences between islands through F_ST_ genetic distances indicates that the population in Isabela is the most divergent of the three, being more divergent to San Cristobal (F_ST_ = 0.207) than to Santa Cruz (F_ST_ = 0.120), coinciding with the geographic distances between the three islands. The genetic distance between Santa Cruz and San Cristobal is considerably lower (F_ST_ = 0.074), suggesting the potential for more widespread gene flow between these two islands throughout the history of *P. guajava* in the archipelago.

**Figure 1.**
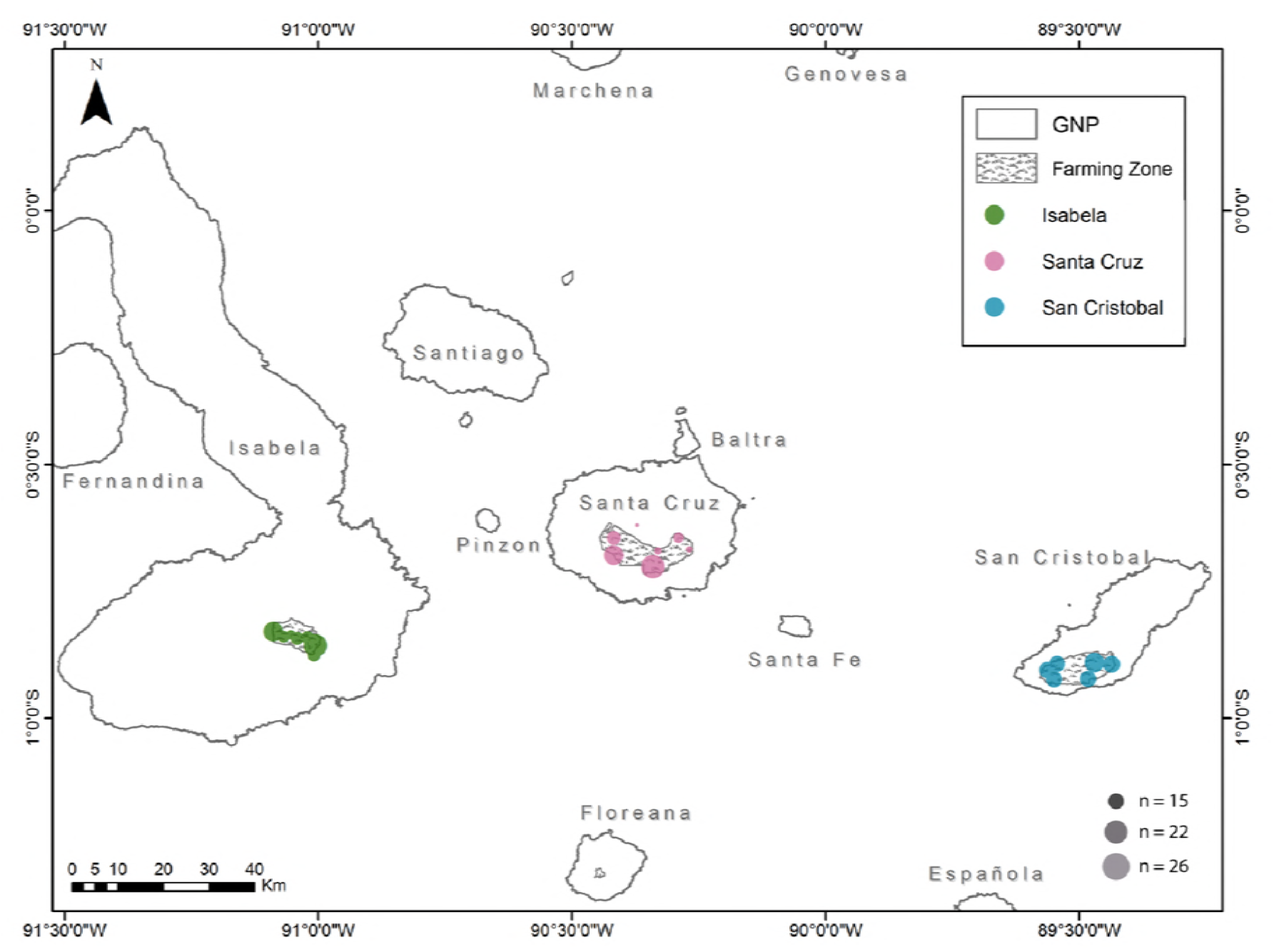
Map representing the sampling sites in three islands from the Galapagos archipelago: Isabela, Santa Cruz and San Cristobal. The diameter of each mark is proportional to the number of samples obtained from each site.

**Figure 2.**
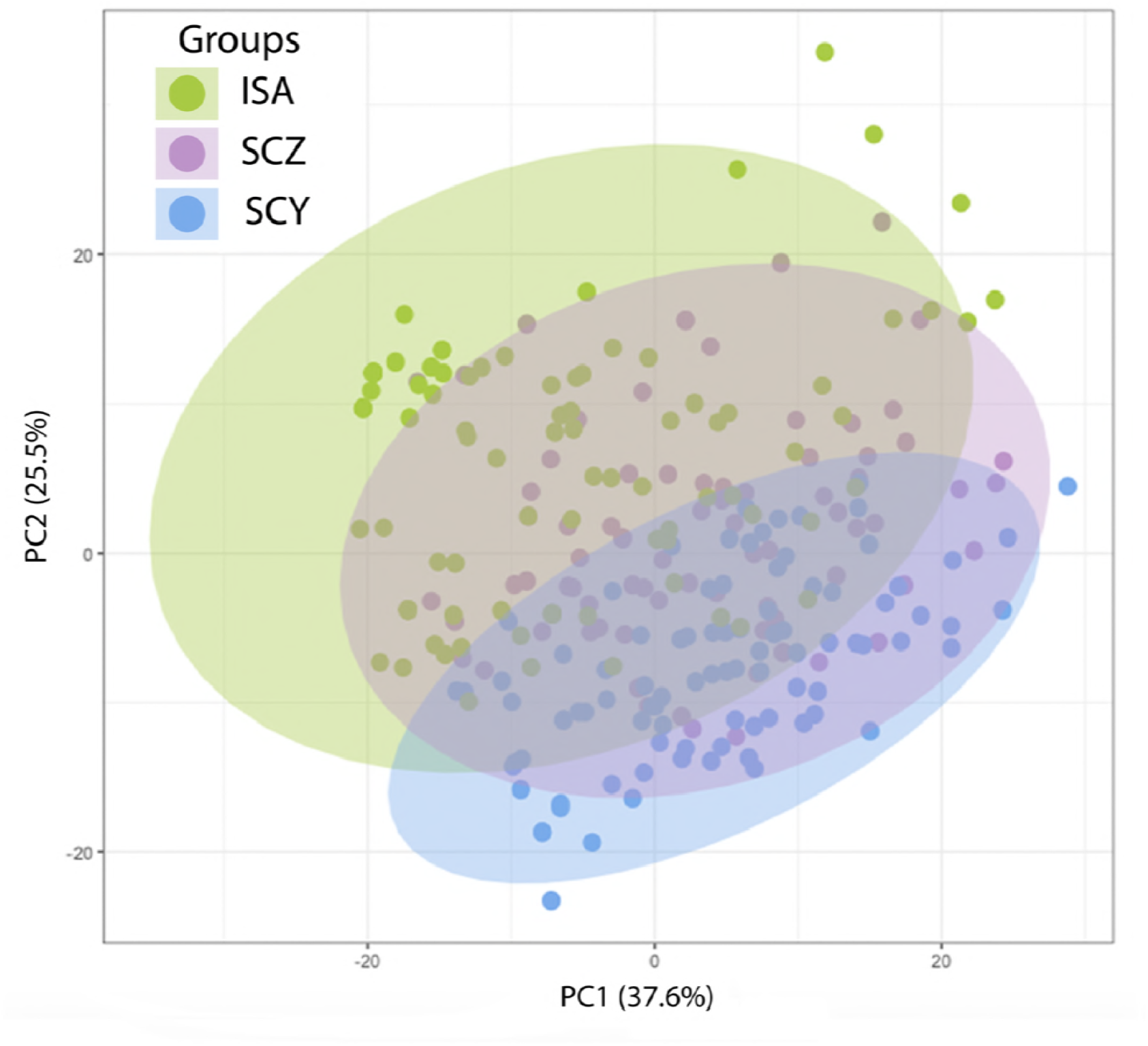
PCoA based on the genetic distances (Euclidian) found between the individuals sampled in the three islands: Isabela (ISA - green), San Cristobal (SCY-blue) and Santa Cruz (SCZ - purple).

**Figure 3.**
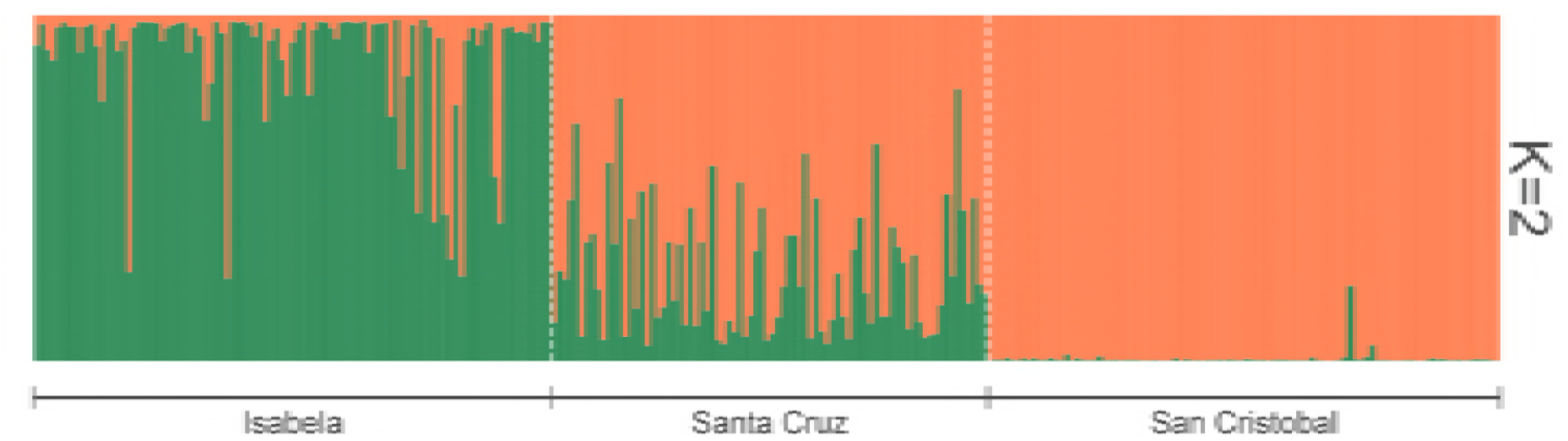
PCoA based on the genetic distances (Euclidian) found between the individuals sampled in the three islands: Isabela (ISA-red dots), San Cristobal (SCY-green dots) and Santa Cruz (SCZ-blue dots).

### Population history of *P. guajava* inferred from genetic marker data

As an invasive species, *P. guajava* has been subject to one or more introduction events in the archipelago, which represent distinct demographic processes that can leave particular genetic footprints in the species’ genome [39]. We tested the possibility of a genetic bottleneck in each island under two distinct SRR mutation models and found significant evidence for a potential population expansion in Isabela (under the SMM model) and a population reduction in San Cristobal (under the TPM model) (Table 3). While these results provide some evidence for specific bottlenecks in specific islands, it also suggests that, under the lack of bottleneck events, there may have been some continuous gene flow between islands (or the archipelago and some continental source population) which reduces the signal of bottleneck events, particularly in Santa Cruz.

**Table 3.** Wilcoxon test results for the support of genetic bottlenecks in the past of the *Psidium guajava* populations of Isabela (ISA), Santa Cruz (SCZ) and San Cristobal (SCY) islands (software BOTTLENECK). Both, the SMM and TPM mutation models implemented in BOTTLENECK were used. One-tailed tests were performed in order to determinate whether the genetic bottleneck occurred because of a heterozygosity (*H*) deficiency (which may happen after a dramatic expansion on the population size) or a *H* excess (which may happen after a dramatic reduction on the population size).

|  | SMM Model |  |  | TPM Model |  |  |
| --- | --- | --- | --- | --- | --- | --- |
|  | ISA | SCZ | SCY | ISA | SCZ | SCY |
| *H* deficiency (Bottleneck due population size expansion) | 0.011* | 0.138 | 0.902 | 0.062 | 0.539 | 0.998 |
| *H* excess (Bottleneck due population size reduction) | 0.992 | 0.884 | 0.125 | 0.949 | 0.500 | 0.004* |
| *H* deficiency or excess (Bottleneck broadly) | 0.021* | 0.275 | 0.250 | 0.123 | 1.000 | 0.008* |

\* indicates *p*-values < 0.05 which support the evidence of a genetic bottleneck.

The population history of *P. guajava* was further investigated through model testing through an Approximate Bayesian Computation (ABC) approach [36] in order to clarify the number of introduction events, the order of these occurrences and the directionality of each event. A total of 14 different scenarios were tested (S1 File), where the two best supported scenarios (posterior probabilities of 0.331 and 0.364) proposed an original introduction event into San Cristobal (Fig. 4). Sub sequentially, the species might have been taken into Isabela first, and later to Santa Cruz (Fig. 4A), or alternatively, the Santa Cruz population might have been formed by introduction events from both San Cristobal and Isabela (Fig. 4B). According the DIYABC estimates, the divergence of the Isabela population from the San Cristobal source (event t2) could date from 455 to 1210 *P. guajava* generations ago, depending on the model. Meanwhile, the split of the Santa Cruz population (event t1) could date from 87 to 275 generations ago. This analysis provides a preliminary hypothesis for the introduction events of *P. guajava* into de Galapagos Islands that fits our current data. It should be noted that we only analyzed individuals from the archipelago and not from any external populations and were therefore unable to determine the source for the current populations in all three islands. It is also still unclear whether an independent introduction event might have occurred for Isabela (event t2(h), Fig. 4C), associated with the high number of exclusive alleles in this population (this event could have occurred from the same continental source population or a different one). This hypothetical scenario could date the separation between the Isabela and San Cristobal populations to before the first introduction of *P. guajava* in the archipelago.

**Figure 4.**
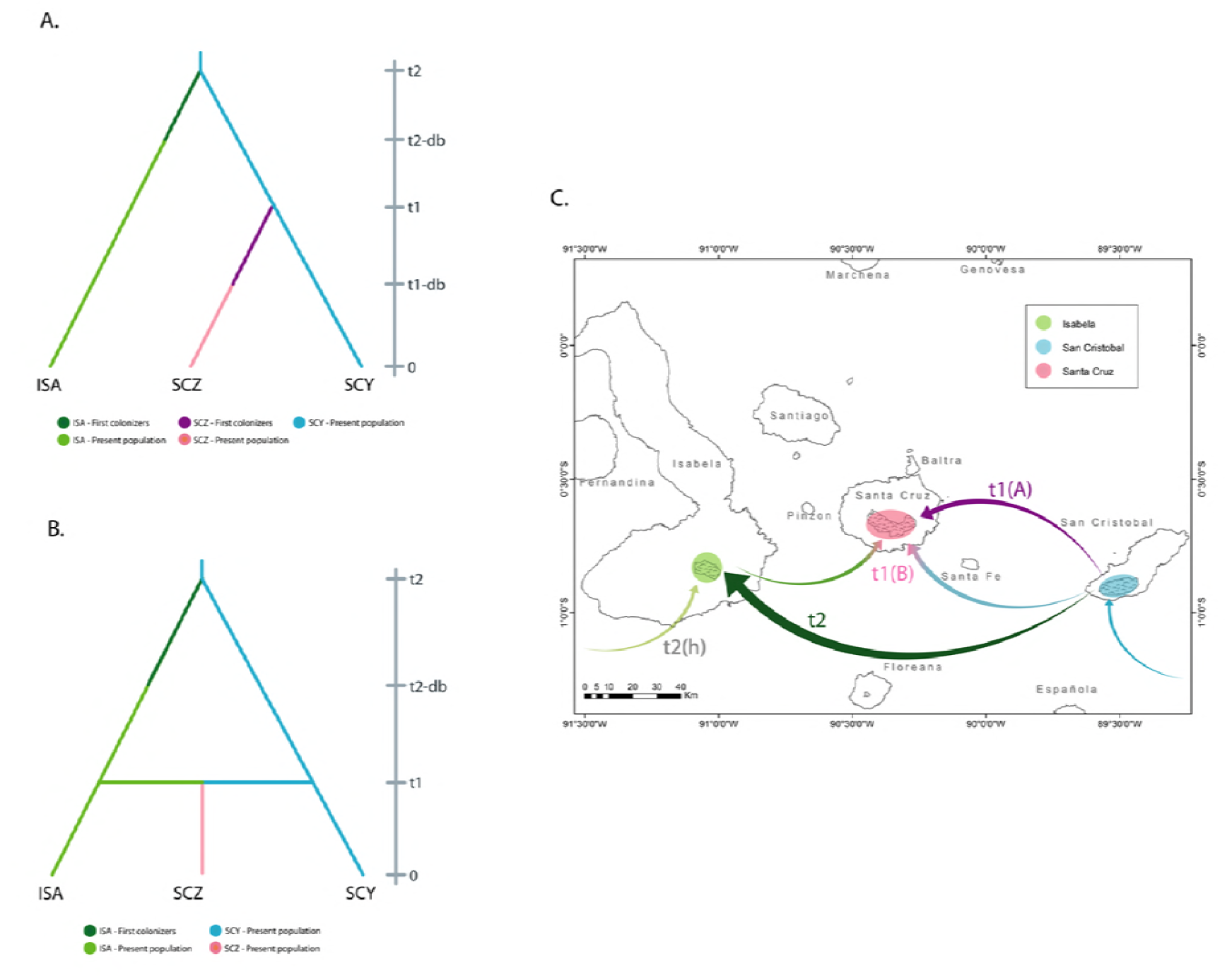
The history of the introduction of *Psidium guajava* in the Galapagos islands. The best models for the introduction history of *P. guajava* were estimated through Approximate Bayesian Computing (ABC) and suggest either a model of independent introductions into Isabela and later Santa Cruz from San Cristobal **(A)**, or an initial introduction into Isabela from San Cristobal followed by introduction events from both islands into Sant Cruz **(B)**. The map **(C)** shows a first introduction into San Cristobal (cyan) from an unknown source (presumably continental South America), which seeded a consecutive introduction into Isabela (t2). The population of Santa Cruz might have been formed by a single introduction from Isabela (t1(A)) or introductions from both San Cristobal and Isabela (t1(B)). It could be proposed that an independent introduction into Isabela also occurred (t2(h)) either before or after the population of San Cristobal.

## Discussion

### Genetic diversity and population structure of *Psidium guajava* in the Galapagos Islands

The overall genetic diversity of *P. guajava* in all three islands (H_E_ = 0.356) is low when compared to the species diversity within its native range, an expected pattern for an insular invasive species [40]. According to research conducted in Brazil [41] and Venezuela [42] with *P. guajava* accessions from germplasm collections, the genetic diversity found in those countries (H_E_ = 0.678 and H_E_ = 0.740 respectively) is considerably higher than that found in the Galapagos *P. guajava* populations. This low genetic diversity in island populations is related to a founder effect, where the first plants that arrive on an island are low in number and are sampled at random from the source population. Because of this, much of the ancestral population’s diversity is left behind, lowering the possibility of obtaining heterozygote individuals [37,43,44]. It is important to consider the size of the colonizing population as well. When the population is small, there will be less diversity compared to the original population [43].

Other plants that are known invaders in island ecosystems, such as *Cortaderia jubata* and *C. selloana* in New Zealand [45] and *Miconia calvescens* in several Pacific islands [46] exhibit a very low genetic diversity, as expected (H_E_ = 0.061, H_E_ = 0.095 and H_E_ = 0.110 respectively). However, other invasive species in island settings, like *Senesio madagascarensis* [47] and *Paraserianthes lophantha* [48] in Hawaii, exhibit much higher genetic diversity levels (H_E_ = 0.790 and H_E_ = 0.600 respectively), argued to be a product of gene flow or multiple introductions [42]. By contrast, the *P. guajava* populations in Galapagos show an expected heterozygosity which is intermediate to these scenarios. This suggests that factors such as multiple introductions, the magnitude of the founder effect, the reproductive system of the species and the geography of the invaded ecosystem [43,49,50] could have affected the genetic diversity of the invasive *P. guajava* populations.

Furthermore, the *P. guajava* populations in different islands from the Galapagos archipelago share some features that provide an insight into the forces that have shaped their current status. The equivalent allelic diversity in all three islands, particularly if rare alleles are excluded, suggests that similar introduction dynamics have shaped these populations, or that a certain degree of homogenization through gene flow between islands has occurred. Upon closer inspection, the significantly higher expected heterozygosity index in Santa Cruz (0.365), when compared to Isabela (0.284) and San Cristobal (0.326), opens the possibility of multiple introductory events that lead to a higher genetic diversity [51]. The low heterozygosity in Isabela and San Cristobal could be explained by a low genetic variation among the first *P. guajava* introduced to these islands, a common occurrence during the colonization of new island ecosystems [40], and a lower number of introduction events.

The degree of differentiation among different islands is noteworthy, as it suggests that gene flow is considerably lower than might be expected for plant populations in different islands. Only 13% of the total differences occur between islands (Table 2), suggesting that gene flow between islands has occurred whether by natural or anthropogenic causes. The major differences of population structure are most likely derived from different introduction events (see below), as the recent history of the species in the archipelago provides with limited opportunity for adaptive processes to produce any notable population stratification [52].

### Factors behind the success of the *Psidium guajava* invasion in the Galapagos

Despite its low genetic diversity, *P. guajava* has proven to be a very aggressive invasive species in the Galapagos [15]. One way to explain this paradox addresses the reproductive biology of *P. guajava*. A proportion of the analyzed individuals showed higher values of the F inbreeding index (S1 Fig), and the mean inbreeding coefficients are high for the three populations (Table 1). These results may be explained, at least partially, by selfing and vegetative reproduction, which occur frequently in *P. guajava* [53]. In the initial stages of its introduction into each island, these reproductive strategies could have allowed this plant to spread rapidly [54]. Then, outcrossing, which occurs with a frequency of 35-40% in *P. guajava* [13], would allow the already widespread population to maintain its genetic diversity and therefore its adaptability and survivability [55,56]. Thus, this capability of combining selfing with outcrossing could be one of the key features that explain why *P. guajava* became a successful, widely distributed and difficult to control species in the local ecosystems.

It should be noted that other non-genetic factors could also explain the success of *P. guajava*. Epigenetics have been shown to play a role in the success of the invasive species *Fallopia japonica*, which despite its low genetic diversity, has high levels of phenotypic variability across populations, which increases its adaptability in new habitats. In this case, the high epigenetic diversity is what explains the fitness of this plant [57]. Phenotypic plasticity could also allow an invasive plant to tolerate a wide range of environmental conditions and to find an ecologic niche in a new habitat [46,58]. *P. guajava* is well adapted to multiple types of soil and luminosity and can also tolerate drought conditions for up to five months [53,59]. Furthermore, the spread of *P. guajava* could have been aided by the human disturbance of native ecosystems. In the Galapagos, abandoned farms became ideal places for this plant to grow and, due to a lack of control, today there are monotypic forests of *P. guajava* on the islands [60]. Human activities and animals such as cattle, pigs and goats remove native vegetation and alter soil, creating space for the invasive species to spread [2,43,61].

The success of the species in the Galapagos and its ability to propagate could also be associated with the lack of population structure within each island. Low levels of genetic differentiation among the regions within a single island are put into evidence by low F_ST_ values between regions (S5 Table – S7 Table), low percentages of genetic differentiation between regions (Table 2) and the absence of a clear population structure within each island (S2 Fig – S4 Fig). A possible explanation for this is an elevated gene flow within islands, a key factor in explaining the spread and success of *P. guajava* as an invasive species in Galapagos. This elevated gene flow could be associated to the dispersion of seeds by birds such as finches, tortoises and certain lizards [62,63], as well as humans and domestic animals [64]. All these factors contribute to the homogenization of allelic frequencies between all the regions within every island. The small size and numerous seeds in each fruit also help *P. guajava* disperse with a great efficiency [65].

### *P. guajava* in the Galapagos: The history of an invasion

The relatively recent colonization of the Galapagos Islands allows us to link the population genetics of the likewise recently introduced *P. guajava* with the well documented historic events that describe the first human settlements on the archipelago. Most of our historical sources are based predominantly on the memories and journals kept by the first settlers and resident sailors of the Galapagos, as well as their descendants’. The information from these sources matches the historical documents obtained from the National Archives, archaeological evidence (e.g. abandoned haciendas and factories from the XIX century in Galapagos) and publications from academic historians [66,67].

Palynological research confirmed that *P. guajava* was present in the highlands of San Cristobal at least since the 1930’s [68]. However, *P. guajava* was documented in historical sources detailing the Galapagos colonization several decades earlier. Early Galapagos settlers wrote about their concern of the invasive species, as they saw it as a potential plague [51,66,69,70]. One of the earliest mentions of *P. guajava* dates from the late XIX century (ca-1889-1890), where it is said that the very first three guava trees on the archipelago were planted by Manuel J. Cobos in his personal garden on San Cristobal Island for ornamental purposes. Cobos was an eminent settler of the Galapagos and was well-known throughout the archipelago. In 1870 he established a massive and profitable sugar cane plantation over the highlands of San Cristobal [66,67]. Unfortunately for the heirs of Cobos, by the 1920’s, the plantation began to be displaced by *P. guajava* monotypic forests [51,66]. In the 1930’s, the plague that had already become a problem in San Cristobal appeared in Floreana island, the first of the Galapagos islands to be colonized by humans [51,70]. Having witnessed the negative effects of this plague in San Cristobal and Floreana, the few permanent settlers of Santa Cruz tried to eradicate any plant or seedling that appeared to be *P. guajava*. However, their attempts failed, and in the 1950’s the plague spread on Santa Cruz [51,66]. *P. guajava* is poorly documented in the Isabela records; according to Lundh (2006) [51], the plague there simply occurred and persists to this day.

#### San Cristobal

The results obtained in this study provide evidence that coincides with the historical sources which suggest that the San Cristobal *P. guajava* population is the origin of the plague that spread over the four inhabited islands of the Galapagos. All but two rare alleles from the San Cristobal population are also present in the Isabela and Santa Cruz populations (Table 1). Furthermore, the STRUCTURE assignment coefficient plot shows that the San Cristobal population lineage (Fig. 3: shown in orange) is present in over 50% of the Santa Cruz population, and to a lesser extent in the Isabela population as well. The PCoA shows the San Cristobal cluster overlapping markedly the Santa Cruz cluster and also some Isabela individuals (Fig. 2). Moreover, this hypothesis is supported by the two best supported scenarios in the ABC analysis. Both of these scenarios show the San Cristobal *P. guajava* population as the ancestral population from which the Isabela and Santa Cruz populations derived (Fig. 4).

The San Cristobal population shows evidence of having gone through a genetic bottleneck which created a transient excess of heterozygosity (Table 3), an event that occurs when the population size is reduced to a few individuals [71,72]. However, it should be considered that, despite Lundh’s [66] affirmation that the San Cristobal plague began with three guava trees planted in Cobos’ garden, San Cristobal received more settlers after Cobos from other islands within the archipelago and the Ecuadorian mainland as well. Therefore, it is possible that some of these new settlers brought new *P. guajava* seeds or seedlings which also contribute to the present-day San Cristobal population. This phenomenon would have increased the genetic diversity of the San Cristobal population to a certain extent but would have not obscured the previously described bottleneck due to a founder effect. The F_IS_ value estimated for this population is markedly lower than the values found for the Santa Cruz and Isabela populations. This lower level of inbreeding might be expected when severe bottlenecks occur, as is the case of the invasive plant *Miconia calvescens*, where very few individuals (or possibly a single individual) planted in Tahiti Island, started an aggressive invasion that spread over several islands of the Pacific [46].

#### Isabela

Despite the fact that *P. guajava* occupies a larger area on Isabela than on any other island within the Galapagos [73], there is practically no information about the history of this invasive species on the island. Lundh [51] simply mentions that the *P. guajava* invasion persists in Isabela and implies that the arrival of several new settlers after the 1950’s triggered the invasion on this island. Nevertheless, both historical events and our results suggest that *P. guajava* could have been present in Isabela several decades before the 1950’s. This island was the third to be permanently colonized by humans, following Floreana and San Cristobal [66]. Antonio Gil was one of the first settlers of Isabela, he arrived in 1897 after having lived in Floreana for four years [67]. Floreana and San Cristobal were the only islands of the Galapagos with permanent and well-established settlements and there was trade between the two islands in the late 19th century [66]. Therefore, it is possible that *P. guajava* was introduced from San Cristobal to Floreana during this period. From here, we could speculate that Antonio Gil, who worked with cattle, might have carried *P. guajava* seeds or fruit to Isabela. This statement is supported by both of the scenarios obtained from our ABC analysis which show that the Isabela population derived from the original population of San Cristobal, before the Santa Cruz population appeared (Fig. 4). The AMOVA results ratify these scenarios and indicate 20% of variation between the populations of San Cristobal and Isabela (Table 2) which suggests important genetic similarities among these populations despite being located on opposite sides of the archipelago.

Both historical and genetic data point to the San Cristobal population as the predecessor of the Isabela population. However, 8 frequent alleles were found in Isabela but not in San Cristobal. Furthermore, several Isabela *P. guajav*a individuals are positioned far away from the San Cristobal cluster in the PCoA (Fig. 2). Also, in the STRUCTURE plot, the Isabela population is mostly clustered in a lineage that is practically absent in San Cristobal (Fig. 3: shown in green). These results suggest that a second independent introduction of *P. guajava* to Isabela from somewhere other than San Cristobal or Floreana is possible, having an important contribution to the present-day Isabela *P.guajava* population and its genetic diversity. This second independent introduction could have occurred either from the same gene pool than the one that seeded San Cristobal or an independent source. In the first case, the ABC divergence time estimates for the San Cristobal and Isabela populations suggests that both islands share a common source population at some date that predates the introduction of *P. guajava* to the Galapagos, presumably a common source continental population. Furthermore, the new contribution of alleles from a second introduction could also have inflated the heterozygosity expected at mutation-drift equilibrium, which is very sensible to a change in the number of alleles [37], thus explaining the heterozygosity deficiency found in the BOTTLENECK analysis.

#### Santa Cruz

Santa Cruz was the last inhabited island of the archipelago to be permanently colonized, and therefore, the last island where *P. guajava* arrived. Settlements began in the 1910’s by former workers of the Cobos plantation in San Cristobal including a foreman, and a single settler from the Ecuadorian mainland. More people arrived in the 1920’s and 1930’s; some of them from the mainland who brought cattle with them, some of them workers from San Cristobal, and several foreigners, especially Norwegians (including sailors from San Cristobal) and Germans [66,67]. *P. guajava* could have been introduced to Santa Cruz in these decades by the abovementioned settlers but, according to historical data, the actual invasion of *P. guajava* in Santa Cruz didn’t begin until the 1950’s [51]. Historical data are in accordance with our hypothesis which suggests that all *P. guajava* populations on the archipelago derived from an ancestral population from San Cristobal. Furthermore, the best supported scenarios from the ABC analysis corroborate that the Santa Cruz population is the last one to derive from the ancestral population from San Cristobal (Fig. 4).

Our results show that the genetic pool of the Santa Cruz *P. guajava* population has an important contribution not only from the San Cristobal population but also from the lineage that was independently introduced in Isabela. This can be observed in the PCoA where the Santa Cruz cluster overlaps with some individuals from Isabela (Fig. 2). The STRUCTURE plot also shows a contribution of the Isabela lineage in the Santa Cruz population (Fig. 3). Finally, one of the best supported scenarios in the ABC analysis (scenario B) confirms that Isabela, along with San Cristobal, contributed to the present-day Santa Cruz population. The multiple origins of the Santa Cruz *P. guajava* population may explain the absence of bottleneck evidence this island (Table 3), since the effects of genetic bottlenecks are softened when multiple introductions occur [48,49]. The admixture of the San Cristobal and Isabela lineages in Santa Cruz, may also help explain the higher genetic diversity and number of lineages (*K*=4, S3 Fig) found.

#### Post-invasion events

The time when the Isabela lineage arrived in Santa Cruz is unclear. During the first half of the XX century, when *P. guajava* was just established in Santa Cruz, Isabela remained isolated due the presence of a penal colony and the lack of a freighter ship serving this island regularly [66]. This fact may explain the observation of private alleles and the conservation of a separate *P. guajava* lineage in Isabela, as unique alleles may have been produced in a short timespan through the naturally high mutation rates of microsatellite regions and an increase in the effective population size of Isabela [74]. However, this does not explain how and when this lineage was established in Santa Cruz. During the 1960’s, the penal colony was removed and more people (not only from the mainland but also from Isabela and San Cristobal) began to move to Santa Cruz, where tourism was being developed the most [66,69,75]. Therefore, it is possible that the arrival of *P. guajava* in Santa Cruz occurred at this time. The important gene flow detected between the populations of San Cristobal and Santa Cruz (F_ST_=0.074) may be dated around these years as well. Meanwhile, the Isabela lineage is almost absent in San Cristobal (Fig. 3), a fact that coincides with the low gene flow detected between these two islands (F_ST_=0.207). It is interesting to note that the population from Santa Cruz, which is located in the center of the archipelago, has gene flow with both the San Cristobal and the Isabela populations. On the other hand, these two populations, located on opposite extremes of the archipelago, show very little gene flow between them. Thus, gene flow between *P. guajava* populations from different islands seems to be correlated with the geographic position of this island. This could reflect more human mobilization between islands that are closer together versus distant islands such as Isabela and San Cristobal. It must be noted that all of the gene flow between *P. guajava* populations located on different islands would be presumably human-mediated, since this invasive species’ natural dispersers cannot travel across the ocean [62,63,64]. Nowadays, this type of gene flow would be completely interrupted due to the strict biosafety regulations in the Galapagos [76] which ban the mobilization of *P. guajava* seedlings, fruit or seeds between islands and between the mainland and the islands (Mónica Ramos, ABG, personal communication).

## Conclusions

Genetic information has increasingly become a powerful tool for the reconstruction of a population’s history and the characterization of its diversity and the forces that shape it [77]. Our current results reveal some of these processes in three *P. guajava* populations from the Galapagos Islands, and provide a valuable insight into the history of the invasive process of this species. However, the addition of new information to the presented framework can provide with finer details of the invasive process and could aid in better understanding the full extent of the risks that *P. guajava* represents to the local ecosystems.

Our reconstruction of the history of the invasion is backed by historical records but was insufficient to explore the source population for the Galapagos *P. guajava* populations. Historical sources pinpoint different provinces in mainland Ecuador as the possible source [52,67,68,75], a fact that could be further corroborated with more widespread sampling of these locations. It is also interesting to consider that Floreana Island might have played an important role in the propagation of *P. guajava* in the archipelago, and it remains the only inhabited island whose guava populations have not been sampled. Further studies will address some of these questions, to continue to elucidate the history of the *P. guajava* invasion.

The purpose of describing an invasive process such as this is to ultimately recognize the extent of the risk it poses to the local flora and fauna, and to assist in the development of effective control and mitigation strategies. The case of *P. guajava* is particularly noteworthy due to the occurrence of a native member of the same genus in the archipelago: *Psidium galapageium*. The tools that have been used to explore the invasion of *P. guajava* could also be used to explore the effects of the co-occurrence of both species in the same spatial context. Firstly, a deeper exploration of the adaptive processes occurring on both species and their signature at the molecular level could allow us to quantify the degree to which both species compete directly with each other, and to account for the toll that *P. guajava* places on its endemic counterpart. Secondly, a complete analysis of the potential for hybridization between these two species can account for the potential noxious effects that *P. guajava* could place on the genetic diversity of the *P. galapageium* populations in the archipelago.

## Acknowledgements

We thank María José Ponce and Sara Ponce for their valuable contributions in the data gathering process during the early stages of research, Leandro Vaca for his assistance in the sampling procedures, as well as Viviana Jaramillo, Venancio Arahana, Juan Francisco Delgado and Liseth Salazar for their contributions to the experimental work. We are grateful for the logistic and administrative assistance of the Galapagos Science Center, a joint research unit from Universidad San Francisco de Quito and the University of North Carolina at Chapel Hill. We also thank Hugo Valdebenito for his comments, support in the field and for sharing information, Marcelo Loyola and Daniel García for their help during the fieldwork, Heinke Jäger (Charles Darwin Foundation) for her suggestions, Maria Mercedes Cobo for proofreading this manuscript and all members from the Plant Biotechnology Laboratory who contributed to this research.

This research was funded by: USFQ GAIAS Grants and USFQ Fondos COCIBA. Genetic data for specimens were obtained under the Genetic Resources Permit Number: MAE-DNB-CM-2016-0041 granted to Universidad San Francisco de Quito by Ministerio del Ambiente Ecuador, in accordance with the Ecuadorian law.

## Supporting Information

**S1 File** Diagrams of the 14 scenarios tested through ABC analysis to infer the history of *Psidium guajava* colonization in the Galapagos Islands. Time is not shown to scale and is measured as number of generations, considering t2>=t1.

Pop1/N1: Present Isabela *P. guajava* population. Pop2/N2: Present Santa Cruz *P. guajava* population. Pop3/N3: Present San Cristobal *P. guajava* population.N1b: Isabela first *P. guajava* colonizers.N2b: Santa Cruz first *P. guajava* colonizers.N3b: San Cristobal first *P. guajava* colonizers.

**S1 Fig.**. Histogram showing the frequency of the *F* (inbreeding coefficient) values observed within the 269 *Psidium guajava* individuals sampled. *F* values were obtained by computing the likelihood function

**S2 Fig.** Results of the Bayesian analysis of population structure (Software STRUCTURE) under the Admixture model, considering only the *Psidium guajava* individuals sampled in Isabela Island. The results are indicated for K = 2, this being the optimum K value (ΔK = 19.62). The values of K correspond to the number of clusters (represented by different colors) in which are grouped the sampled individuals. White dotted lines separate different regions.

**S3 Fig.** Results of the Bayesian analysis of population structure (Software STRUCTURE) under the Admixture model, considering only the *Psidium guajava* individuals sampled in Santa Cruz Island. The results are indicated for K = 4, this being the optimum K value (ΔK = 39.57). The values of K correspond to the number of clusters (represented by different colors) in which are grouped the sampled individuals. White dotted lines separate different regions (GR=Granillo Rojo).

**S4 Fig.** Results of the Bayesian analysis of population structure (Software STRUCTURE) under the Admixture model, considering only the *Psidium guajava* individuals sampled in San Cristobal Island. The results are indicated for K = 2, this being the optimum K value (ΔK = 1.83). The values of K correspond to the number of clusters (represented by different colors) in which are grouped the sampled individuals. White dotted lines separate different regions.

**S1 Table.** Percentage of missing data for the three islands, missing data per primer and the total missing data for all individuals (269).

**S2 Table.** Populations in which a significant linkage disequilibrium (LD) between pairs of loci was found, after Bonferroni Correction. The names of the linked loci appear in rows and columns, whereas population names appear as entries in the table. ISA= Isabela population; SCY= San Cristobal population. No LD was found in Santa Cruz.

**S3 Table.** Results of the Hardy-Weinberg Equilibrium (HWE) test for each loci within the three island populations (Isabela, Santa Cruz, San Cristobal). Results shown correspond to those after Bonferroni correction.

**S4 Table.** Results of the analysis of molecular variance (AMOVA) performed within the *Psidium guajava* populations from Isabela, Santa Cruz and San Cristobal islands, individually. Missing data was ignored for this analysis.

**S5 Table.** Pairwise F_ST_ values between all the Isabela Island regions in which *Psidium guajava* individuals were sampled.

**S6 Table.** Pairwise F_ST_ values between all the Santa Cruz Island regions in which *Psidium guajava* individuals were sampled.

**S7 Table.** Pairwise F_ST_ values between all the San Cristobal Island regions in which *Psidium guajava* individuals were sampled.

## References

1. Cronk QBC, Fuller JL. Plant Invaders: The Threat to Natural Ecosystems Abingdon: Earthscan Publications Ltd.; 2001.

2. Sakai AK, Allendorf FW, Holt JS, Lodge DM, Molofsky J, With KA, et al. The Population Biology of Invasive Species. Annual Review of Ecology and Systematics. 2001;32: 305–333.

3. Weidenhamer JD, Callaway RM. Direct and Indirect Effects of Invasive Plants on Soil Chemistry and Ecosystem Function. Journal of Chemical Ecology. 2010;36: 59–69.

4. Vila M, Espinar JL, Hejda M, Hulme PE, Jarosik V, Maron JL, et al. Ecological impacts of invasive alien plants: a meta-analysis of their effects on species, communities and ecosystems. Ecology Letters. 2011;14: 702–708.

5. Rentería JL, Gardener MR, Panetta FD, Atkinson R, Crawley MJ. Possible impacts of the invasive plant *Rubus niveus* on the Native Vegetation of the Scalesia Forest in the Galapagos Islands. PLoS ONE. 2012; 7(10): e48106.

6. Mauchamp A. Threats from Alien Plant Species in the Galápagos Islands. Society for Conservation Biology. 1997;11(1): 260–263.

7. Mooney HA, Cleland EE. The evolutionary impact of invasive species. Proceedings of the National Academy of Sciences. 2001;98(10): 5446–5451.

8. Bensted-Smith R. A Biodiversity Vision for the Galapagos Islands. Puerto Ayora: Charles Darwin Foundation, World Wildlife Foundation; 2002.

9. Toral-Franda MV, Causton CE, Jager H, Trueman M, Izurieta JC, Araujo E, et al. Alien species pathways to the Galapagos Islands, Ecuador. PLoS ONE. 2017;12(9): e0184379.

10. Tye A. Can we infer island introduction and naturalization rates from inventory data? Evidence from introduced plants in Galapagos. Biological Invasions. 2006; 8: 201–215.

11. Charles Darwin Foundation. Research. 2018 [cited 2018 March 22]. Available from: https://www.darwinfoundation.org/en/research.

12. Walsh SJ, McCleary AL, Mena CF, Shao Y, Tuttle JP, González A, et al. QuickBird and Hyperion data analysis of an invasive plant species in the Galapagos Islands of Ecuador: Implications for control and land use management. Remote Sensing of Environment. 2008; 112: 1927–1941.

13. Sitther V, Zhang D, Harris DL, Yadav AK, Zee FT, Meinhardt LW, et al. Genetic characterization of guava (*Psidium guajava* L.) germplasm in the United States using microsatellite markers. Genetic Resources Crop Evolution. 2014;61: 829–839.

14. Velasco M. Percepciones de la población de Galápagos sobre las especies introducidas y Sistema de Inspección y Cuarentena para Galápagos (SICGAL). Darwin FC, editor. Quito; 2002.

15. Tye A, Atkinson R, Carrión V. Increase in the number of introduced plant species in Galapagos. In: DPNG, GCREG, FCD, GC. Galapagos Report 2006-2007. Puerto Ayora: DPNG; 2007. pp. 133–135.

16. Torres MdL, Mena CF. Understanding Invasive Species in the Galapagos Islands: Springer International Publishing AG 2018; 2018.

17. Levin DA, Francisco-Ortega J, Jansen RK. Hybridization and the Extinction of Rare Plant Species. Conservation Biology. 1996;10(1): 10–16.

18. López-Caamal A, Cano-Santana Z, Jiménez-Ramírez J, Ramírez-Rodríguez R, Tovar-Sánchez E. Is the insular endemic *Psidium socorrense* (Myrtaceae) at risk of extinction through hybridization? Plant Systematics and Evolution. 2014;300(9): 1959–1972.

19. Shaik R, Zhu, Clements DR, Weston LA. Understanding invasion history and predicting invasive niches using genetic sequencing technology in Australia: case studies from Cucurbitaceae and Boraginaceae. Conservation Physiology. 2016;4(1): cow030.

20. Saghai-Maroof MA, Soliman KM, Jorgensen RA, Allard RW. Ribosomal DNA spacer-length polymorphisms in barley: mendelian inheritance, chromosomal location, and population dynamics. Proceedings of the National Academy of Sciences of the United States of America. 1984;81(24): 8014–8018.

21. Risterucci A, Duval MF, Rohde W, Billotte N. Isolation and characterization of microsatellite loci from *Psidium guajava* L. Molecular Ecology Notes. 2005;5: 745–748.

22. Blacket MJ, Robin C, Good RT, Lee SF, Miller AD. Universal primers for fluorescent labelling of PCR fragments--an efficient and cost-effective approach to genotyping by fluorescence. Molecular Ecology Resources. 2012;12(3): 456–463.

23. Goudet J. hierfstat, a package for R to compute and test hierarchical F-statistics. Molecular Ecology Notes. 2005;5(1): 184–186.

24. Team RC. R: A language and environment for statistical computing. Vienna, Austria. URL http://www.R-project.org. 2015.

25. Kamvar ZN, Tabima JF, Grünwald NJ. Poppr: an R package for genetic analysis of populations with clonal, partially clonal, and/or sexual reproduction. PeerJ. 2014;2: e281.

26. Excoffier L, Lischer HEL. Arlequin suite ver 3.5: A new series of programs to perform population genetics analyses under Linux and Windows. Molecular Ecology Resources. 2010;10: 564–567.

27. Jombart T, Ahmed I. adegenet 1.3-1: new tools for the analysis of genome-wide SNP data. Bioinformatics. 2011;27(21): 3070–3071. pmid:21926124

28. Paradis E. pegas: an R package for population genetics with an integrated—modular approach. Bioinformatics. 2010;26(3): 419–420. pmid:20080509.

29. Weir BS, Cockerham CC. Estimating F-statistics for the analysis of population structure. Evolution. 1984;38(6): 1358–1370.

30. Beck MW. ggord: Ordination Plots with ggplot2. 2017 [cited 4 April 2018]. In: zenodo [Internet]. fawda123/ggord: v1.1.0. Available from: https://zenodo.org/badge/latestdoi/35334615

31. Pritchard JK, Stephens M, Donelly P. Inference of Population Structure Using Multilocus Genotype Data. Genetics. 2000;155: 945–959.

32. Evanno G, Regnaut S, Goudet J. Detecting the number of clusters of individuals using the software STRUCTURE: a simulation study. Molecular Ecology. 2005;14(8): 2611–2620.

33. Earl DA, von Holdt BM. STRUCTURE HARVESTER: a website and program for visualizing STRUCTURE output and implementing the Evanno method. Conservation Genetics Resources. 2012;4(2): 359–361.

34. Jakobsson M, Rosenberg NA. CLUMPP: a cluster matching and permutation program for dealing with label switching and multimodality in analysis of population structure. Bioinformatics. 2007;23(14): 1801–1806.

35. Francis RM. pophelper: an R package and web app to analyse and visualize population structure. Mol Ecol Resour. 2017;17(1): 27–32.

36. Cornuet JM, Pudlo P, Veyssier J, Dehne-Garcia A, Gautier M, Leblois R, et al. DIYABC 2.0: a software to make approximate Bayesian computation inferences about population history using single nucleotide polymorphism, DNA sequence and microsatellite data. Bioinformatics. 2014;30(8): 1187–1189.

37. Piry S, Luikart G, Cornuet JM. BOTTLENECK: a computer program for detecting recent reductions in the effective size using allele frequency data. Journal of Heredity. 1999;90(4): 502–503.

38. Marshall DR, Brown AH. Optimum sampling strategies in genetic conservation. In: Frankel OH, Hawkes JG, editors. Crop Genetic Resources for Today and Tomorrow. Cambridge: Cambridge University Press; 1975. pp. 53–80.

39. Luikart G, Allendorf FW, Cornuet JM, Sherwin WB. Distortion of allele frequency distributions provides a test for recent population bottlenecks. Journal of Heredity. 1998;89(3): 238–247.

40. Lee C. Evolutionary genetics of invasive species. Trends Ecol Evol. 2002; 17(8): 386–391.

41. Da Costa SR, Santos CaF. Allelic database and divergence among *Psidium* accessions by using microsatellite markers. Genetics and Molecular Research: GMR. 2013;12(4): 6802–6812.

42. Aranguren Y, Briceño A, Fermin G. Assessment of the Variability of Venezuelan Guava Landraces by Microsatellites. Acta Hort. 2010;849(16): 147–154.

43. Stuessy TF, Takayama K, López-Sepúlveda P, Crawford DJ. Interpretation of patterns of genetic variation in endemic plant species of oceanic islands. Botanical Journal of the Linnean Society. 2014;174(3): 276–288.

44. Frankham R. Do island populations have less genetic variation than mainland populations? Heredity. 1997;78(3): 311–327.

45. Houliston G, Goeke D. *Cortaderia spp*. In New Zealand: Patterns of genetic variation in two widespread invasive species. New Zealand Journal of Ecology. 2017;41: 107–112.

46. Le Roux J, Wieczorek A, Meyer JY. Genetic diversity and structure of the invasive tree *Miconia calvescens* in Pacific islands. Diversity and Distributions. 2008;14: 935–948.

47. Le Roux J, Wieczorek A, Carol T, Vorsino A. Disentangling the dynamics of invasive fireweed (*Senecio madagascariensis* Poir. species complex) in the Hawaiian Islands. Biol Invasions. 2010;12(7): 2251–2264.

48. Thompson GD, Richardson DM, Wilson JRU, Bellstedt DU, Roux JJL. Genetic diversity and structure of the globally invasive tree, P*araserianthes lophantha* subspecies *lophantha*, suggest an introduction history characterised by varying propagule pressure. Tree Genetics & Genomes. 2016. doi: 10.1007/s11295-016-0984-0

49. Barrett SC, Husband BC. In: Brown AH, Clegg MT, Kahler AL, Weir BS. Plant population genetics, breeding, and genetic resources. Sunderland: Sinauer Associates Inc.; 1990. pp. 254–277.

50. Hagenblad J, Hülskötter J, Acharya K, Brunet J, Chabrerie O, Cousins S, et al. Low genetic diversity despite multiple introductions of the invasive plant species *Impatiens glandulifera* in Europe. BMC Genet. 2015. doi: 10.1186/s12863-015-0242-8

51. Lundh JP. The farm area and cultivated plants on Santa Cruz, 1932-1965, with remarks on other parts of Galapagos. Galapagos Research. 2006;64: 12–25.

52. Barton N. Understanding Adaptation in Large Populations. PLoS Genet. 2010;6(6): e1000987.

53. Binggeli P, Hall JB, Healey JR. A review of invasive woody plants in the tropics. 1st ed. Bangor: University of Wales; 1998.

54. Baker HG. Self-compatibility and establishment after ‘long-distance’ dispersal. Evolution. 1955;9: 347–349.

55. Carlquist S. Island biology. 1st ed. New York: Columbia University Press; 1974.

56. Crawford DJ, Lowrey TK, Anderson GJ, Bernardello G, Santos-Guerra A, Stuessy TF. Genetic diversity in Asteraceae endemic to oceanic islands: Baker’s Law and polyploidy. In: Funk VA, Susanna A, Stuessy TF, Bayer RJ. Systematics, evolution, and biogeography of Compositae. Vienna: IAPT; 2009. pp. 139–151.

57. Richards C, Schrey A, Pigliucci M. Invasion of diverse habitats by few Japanese knotweed genotypes is correlated with epigenetic differentiation. Ecology Letters. 2012;9(10): 16–26.

58. Sol D. Claves del éxito de las especies invasoras. Ambiente. 2014;109: 4–13.

59. Somarriba E. Effects of livestock on seed germination of guava (*Psidium guajava* L.). Agroforestry Systems. 1986;4(3): 233–238.

60. Chiriboga R, Fonseca B, Maignan S. Desarrollo de políticas y estrategias de manejo del sector Agropecuario y su relación con las especies introducidas en la Provincia de Galápagos. In: Proyecto ECU/00/G31. Diagnóstico agrario en las islas Galápagos. Quito: SIPAE-PNUD-INGALA; 2006.

61. Bramwell D. Plants and Islands. 1st ed. London: Academic Press; 1979.

62. Heleno RH, Olesen JM, Nogales M, Vargas P, Traveset A. Seed dispersal networks in the Galápagos and the consequences of alien plant invasions. Proc R Soc B. 2013. doi: 10.1098/rspb.2012.2112.

63. Blake S, Wikelski M, Cabrera F, Guezou A, Silva M, Sadeghayobi E, et al. Seed dispersal by Galapagos tortoises. Journal of Biogeography. 2012;39(11): 1961–1972.

64. Herrera XM. Posibles dispersores de *Psidium guajava* en la Isla San Cristóbal, Galápagos – Ecuador. B.Sc. Thesis, Universidad San Francisco de Quito. 2013. Available from: http://repositorio.usfq.edu.ec/handle/23000/2777

65. Henderson, S. Species profile: *Psidium guajava*; 2010 [cited 2017 Sept 10]. Database: Global Invasive Species Database (GISD) [Internet]. Available from: http://www.iucngisd.org/gisd/species.php?sc=211

66. Lundh JP. Galápagos: A Brief History; 2004 [cited 2017 Sept 20]. Repository: Human and Cartographic History of the Galápagos Islands [Internet]. Available from: http://www.galapagos.to/TEXTS/LUNDH1-1.php

67. Latorre O. Galapagos: Los primeros habitantes de algunas islas. Noticias de Galápagos. 1997;56–57: 62–66.

68. Restrepo A, Colinvaux P, Bush M, Correa-Metrio A, Conroy J, Gardener MR, et al. Impacts of climate variability and human colonization on the vegetation of the Galápagos Islands. Ecology. 2012;93(8): 1853–1866.

69. Lundh JP. The Last Days of a Paradise; 2006 [cited 2018 May 8]. Repository: Human and Cartographic History of the Galápagos Islands [Internet]. Available from: http://www.galapagos.to/TEXTS/LUNDH3-1.php

70. Strauch D. Satan Came to Eden; 1935 [cited 2018 May 7]. Repository: Human and Cartographic History of the Galápagos Islands [Internet]. Available from: http://www.galapagos.to/TEXTS/STRAUCH-1.php

71. Estoup A, Angers B. Microsatellites and Minisatellites. In: Carvalho G, editor. Advances in Molecular Ecology. Amsterdam: IOS Press; 1998. pp. 55–79.

72. Luikart G, Cornuet JM. Empirical Evaluation of a Test for Identifying Recently Bottlenecked Populations from Allele Frequency Data. Conservation Biology. 1998;12(1): 228–237.

73. Rivas-Torres GF, Benítez FL, Rueda D, Sevilla C, Mena CF. A methodology for mapping native and invasive vegetation coverage in archipelagos. Progress in Physical Geography. 2018;42(1): 83–111.

74. Marriage TN, Hudman S, Mort ME, Orive ME, Shaw RG, Kelly JK. Direct estimation of the mutation rate at dinucleotide microsatellite loci in *Arabidopsis thaliana* (Brassicaceae). Heredity. 2009;103(4): 310–317.

75. Granda M, Chóez G. Sistemas Humanos: Población y Migración en Galápagos. In: DPNG, GCREG, FCD, GC. Informe Galápagos 2011-2012. Puerto Ayora: DPNG; 2013. pp. 44–51.

76. Registro Oficial Suplemento 989 de 21-abr.-2017. Decreto Ejecutivo 1363: No. 1363.

77. Cristescu ME. Genetic reconstructions of invasion history. Mol Ecol. 2015;24: 2212–2225.

